# MYORG and STRADB as Activity-Dependent Therapeutic Targets for Frailty Prevention: Discovery and Cross-Cohort Validation in Aging Skeletal Muscle

**DOI:** 10.64898/2026.04.18.719222

**Authors:** Rangaprasad Sarangarajan, Kastoori Iyengar

## Abstract

**Background:** Skeletal muscle aging exhibits substantial heterogeneity, with some individuals maintaining robust function into advanced age while others develop sarcopenia and frailty. Whether molecular signatures distinguishing these trajectories reflect biological aging or modifiable factors, such as physical activity, remains unclear.

**Methods:** An integrated discovery-validation study was conducted on skeletal muscle transcriptomes. Discovery analysis used the GSE144304 dataset comprising vastus lateralis biopsies from young adults (n=26, aged 18-30 years), fit elderly (n=30, aged 65-80 years with preserved function), and frail elderly (n=24, aged 65-80 years stratified by grip strength). Top 10 most significantly altered genes were validated across five independent transcriptomic studies (n=184 total) strategically selected to represent distinct activity contexts: activity-controlled aging, sedentary aging, mixed-activity aging, disease-impaired aging, and exercise intervention. Expression of two established atrogenes were examined (FBXO32/Atrogin-1 and TRIM63/MuRF-1) as benchmarks.

**Results:** Discovery analysis identified 10 genes with profound age-related changes (adjusted p < 10⁻²¹, |log₂FC| > 1.3). Cross-dataset validation revealed striking activity-dependence: genes downregulated with aging in sedentary populations (MYORG, STRADB) showed maintained or increased expression in active elderly individuals (80% validation rate, r = 0.75-0.82 with activity level). In contrast, established atrogenes showed poor replication (25-50%) and context-dependent patterns. C4ORF54 expression strongly correlated with grip strength (r = 0.68, p < 0.001), with age effects disappearing after phenotype adjustment, indicating purely phenotype-mediated expression. Critically, sedentary versus active aging datasets showed opposing transcriptional patterns (r = −0.68), demonstrating that activity confounds conventional age-based signatures.

**Conclusions:** Molecular signatures distinguishing fit from frail aging predominantly reflect physical activity levels rather than inevitable biological processes. MYORG and STRADB emerge as activity-responsive biomarkers of muscle health, while C4ORF54 serves as an indicator of functional capacity. These findings challenge conventional atrogene paradigms and suggest that exercise-responsive AMPK signaling pathways represent immediately translatable therapeutic targets for preserving muscle function in older adults.

## INTRODUCTION

Sarcopenia, the progressive loss of muscle mass, strength, and function with aging, is a major contributor to frailty, disability, and mortality in older adults(Abellan Van Kan, 2009; Degens, n.d.; Newman, 2023). Yet its trajectory varies dramatically between individuals, wherein some elderly individuals maintain robust muscle function while others of identical chronological age develop severe impairment(Faulkner et al., 2007; Granic et al., 2026). This heterogeneity implies that muscle aging is shaped by modifiable factors rather than by immutable biology.

Transcriptomic profiling has been used to discover characteristic gene expression changes in aging muscle, typically via cross-sectional comparisons between young and old adults, including downregulation of contractile, metabolic, and mitochondrial genes alongside upregulation of inflammatory, extracellular matrix, and stress-response pathways(Tieland et al., 2018; Tumasian et al., 2021; Welle et al., 2003; Wilkinson et al., 2018). Cross-sectional studies are confounded by the fact that physical activity declines substantially with age, independent of functional capacity. Since disuse alone profoundly alters muscle gene expression within weeks, the question arises of whether the observed transcriptomic changes reflect chronological aging itself (primary aging) or reduced physical activity (secondary aging). Primary aging signatures would indicate targets for pharmacological intervention; secondary aging signatures would point to preventable changes that can be addressed through lifestyle modification. The molecular underpinnings connecting the two remain a work in progress.

The ubiquitin ligases FBXO32 (Atrogin-1) and TRIM63 (MuRF-1) illustrate the problem. These canonical atrogenes are rapidly upregulated by disuse, denervation, and disease, and their reported upregulation in aging muscle is widely cited as evidence that increased protein degradation drives sarcopenia, influencing therapeutic efforts targeting proteasome inhibition(de Souza et al., 2025; Fernando et al., 2019; Sartori et al., 2021; Taillandier & Polge, 2019). Yet the evidence in the literature is inconsistent. While atrogenes are classic markers of muscle wasting, their role in aging-related declines in skeletal muscle function remains unclear. Furthermore, not all older individuals experience the same degree of muscle decline. While some elderly maintain relatively preserved muscle function (“fit aging”), others develop marked weakness and functional impairment (“frailty”)(Manca, A. et al., 2024; Navarrete-Villanueva et al., 2021; Rockwood et al., 2004). The molecular determinants distinguishing these trajectories remain incompletely understood.

To address this critical gap, we conducted a comprehensive multi-cohort validation study to distinguish activity-dependent from activity-independent transcriptional changes in aging muscle. We first identified the most significantly altered genes in a discovery cohort with detailed functional phenotyping (grip strength), and then systematically validated these findings across 5 independent datasets spanning diverse activity levels, from sedentary to lifelong-trained individuals. By comparing expression patterns across activity-stratified cohorts, we aimed to determine: (1) which “aging genes” are activity-dependent and potentially reversible, (2) whether established atrogenes consistently mark sarcopenia across contexts, and (3) whether specific genes track functional capacity independent of chronological age.

Our findings reveal that secondary aging, mediated primarily through declining physical activity, dominates the skeletal muscle transcriptome signature previously attributed, in part, to chronological aging. This paradigm shift has immediate implications for biomarker development, therapeutic target selection, and preventive interventions for sarcopenia.

## METHODS

### 1. Discovery Analysis

#### Dataset, Study Population, Study Design

This was a secondary analysis and multi-cohort validation study. A query of the GEO database for “human, aging, vastus lateralis” returned 146 studies, sorted based on highest to lowest sample size. The GSE144304 dataset was identified and used for analysis due to the availability of frailty assessment (grip strength, 400m walk test, gait [SPPB score] and Fried frailty score) associated with each individual skeletal muscle sample transcriptomic data from young adults (n=26, age 20-30 years), older fit individuals (n=30, age ≥75 years with preserved physical function), and older frail individuals (n=24, age ≥75 years). Details on the study design, population, phenotypic characteristics, and gene expression profiling methodology for the GSE144304 dataset have been previously published.

#### Differential Expression Analysis

Differential gene expression (DEGs) analysis was performed using the NCBI GEO2R, a web-based tool that implements the limma Bioconductor package for differential analysis, using NCBI-computed raw count matrices as input(Clough et al., 2024). The samples from the list were manually assigned to the following groups: (1) Young, (2) Old (Fit + Frail), (3) Old-Fit and (3) Old-Frail, based on sample metadata. The tool automatically generated an R design matrix to enable differential expression analysis (DEA) between the groups (Young vs. Old; Young vs. Old-Fit; Young vs. Old-Frail; Old-Fit vs. Old-Frail). Genes were considered significantly differentially expressed at adjusted p-value <0.05 (Benjamini-Hochberg correction). Log2 fold change was set to 0 to determine the true positive or negative fold change for genes. To focus on high-confidence, biologically relevant changes, a robust expression threshold requiring baseMean ≥ 500 (average normalized count across all samples, corresponding to approximately 10 transcripts per million [TPM]) was applied. This stringent filter excludes lowly expressed genes prone to technical noise while retaining genes with substantive transcriptional activity, reducing the multiple-testing burden and ensuring that detected changes reflect biologically significant gene products. Significantly altered genes were classified as either ‘declining with age’ (higher expression in young, positive log2FC) or ‘increasing with age’ (higher expression in old, negative log2FC). For the Fit vs Frail comparison, genes were similarly classified as ‘fit-specific’ (higher in fit elderly) or ‘frail-specific’ (higher in frail elderly).

The primary comparisons were conducted to address distinct scientific questions. First, the comparison of young adults to all elderly individuals (fit and frail combined) was used to identify overall age-associated transcriptomic changes. Second, young adults were compared separately to old-fit to characterize changes in successful aging. Third, young adults were compared to the old-frail to characterize changes in unsuccessful aging with frailty development. An additional direct comparison of fit versus frail elderly was performed to identify genes specifically distinguishing aging trajectories. This comparison was particularly valuable because both groups were matched on chronological age, isolating functional differences from age effects.

For each comparison, linear models were fit using the limma framework with empirical Bayes shrinkage of gene-wise variances toward a common value. This approach borrows information across genes to improve statistical power, particularly valuable for genes with small sample sizes relative to biological variability. Moderated t-statistics were calculated for each gene, and p-values were adjusted for multiple testing using the Benjamini-Hochberg false discovery rate procedure. Genes showing adjusted p-values ≤ 0.05 were considered statistically significant.

#### Gene Prioritization for Validation

Based on the differential expression results, we prioritized genes for validation using multiple criteria. Statistical significance was the primary criterion, with genes ranked by adjusted p-value. Next, the focus was on the top 10 most significant genes that met the baseMean threshold. Additional considerations included biological plausibility (genes with known or suspected roles in muscle function), consistency across comparisons (genes showing similar directional changes in multiple comparisons), and effect size (genes showing substantial fold changes).

#### Functional Annotation

Genes were annotated using information from the NCBI Gene database, including official symbols, full names, chromosomal locations, and functional descriptions. Gene Ontology (GO) terms were retrieved to identify biological processes, molecular functions, and cellular components associated with each gene. KEGG pathway assignments identified signaling and metabolic pathways represented among differentially expressed genes. Pathway enrichment analysis was performed using multiple complementary tools, including DAVID (Database for Annotation, Visualization and Integrated Discovery) and standard GO enrichment procedures. For each pathway or GO term, we calculated enrichment statistics comparing the proportion of differentially expressed genes in that category versus the proportion expected by chance. Statistical significance was assessed using hypergeometric tests with false discovery rate correction. Pathways showing FDR-corrected p-values ≤ 0.05 were considered significantly enriched.

### 2. Validation Analysis Dataset Selection

The validation phase required reviewing GEO datasets to identify specific studies that represent distinct activity contexts, enabling assessment of activity dependence. We identified five independent studies that met stringent criteria, including the use of vastus lateralis muscle (matching the discovery tissue), high-quality RNA-sequencing data, adequate sample sizes, and clear documentation of age groups or intervention protocols.

The first validation dataset (GSE159217) represented activity-controlled aging. This study recruited young males (n=20, aged 19-25 years) and older males (n=18, aged 65-71 years) who were specifically matched for physical activity levels (Lagerwaard et al., 2021). Activity matching was verified through both self-reported questionnaires and objective accelerometry measurements over multiple days. Both groups met physical activity guidelines, ensuring that observed transcriptomic differences reflected chronological aging independent of activity-related changes. This design isolates primary aging effects by controlling for secondary aging due to activity decline.

The second validation dataset (GSE175495) represented sedentary aging without activity control. Young adults (n=12, aged 20-35 years) were compared to older adults (n=12, aged 60-85 years) following overnight fasting and 48-hour restriction from strenuous physical activity (Trim et al., 2022). No attempt was made to match activity levels, representing typical aging studies where elderly individuals likely have lower habitual activity than young adults. This design captures the combination of primary and secondary aging effects typically seen in conventional studies.

The third validation dataset (GSE164471) represented mixed-activity aging in a population-based sample. This study enrolled 53 healthy adults spanning a continuous age range from 22 to 83 years without activity matching or intervention (Tumasian et al., 2021). This heterogeneous sample represents real-world population variability in both age and activity levels, enabling assessment of population-level aging patterns.

The fourth validation dataset (GSE242202) represented disease-impaired aging with enforced reductions in activity. This study compared young healthy adults (n=15, aged 26-45 years) to older adults with knee osteoarthritis (n=37, aged 65-83 years) (Kurochkina et al., 2024).

Osteoarthritis causes pain and functional limitations, substantially reducing physical activity independent of motivation. This represents aging with both disease burden and disease-imposed inactivity, enabling assessment of how chronic inflammation and enforced disuse affect gene expression.

The fifth validation dataset (GSE151066) represented an acute exercise intervention in previously sedentary elderly (Rubenstein et al., 2022). This within-subject study enrolled 9 older adults (aged 65-80 years) who were sedentary at baseline. Muscle biopsies were obtained before exercise, immediately after acute endurance exercise, and three hours post-exercise. This design provides mechanistic evidence for activity-dependent gene regulation and captures the acute response to mechanical and metabolic stimulation.

All validation datasets utilized the same tissue (vastus lateralis) and sequencing technology (RNA-seq) as the discovery study, minimizing technical confounding.

#### Data Extraction and Processing

For datasets with published differential expression results (GSE159217, GSE164471, GSE151066), we extracted log2 fold changes and adjusted p-values for our target genes directly from GEO2R analysis of raw data. For datasets providing only normalized expression values (GSE175495, GSE242202), we performed independent statistical analysis. For these datasets, we downloaded normalized gene expression matrices from GEO. We identified samples belonging to the comparison groups using metadata files. For each target gene, we calculated the mean expression in each group and performed unpaired two-sample t-tests comparing groups. P-values were adjusted for multiple testing across all tested genes using the Benjamini-Hochberg procedure. Genes showing p-values ≤ 0.05 were considered statistically significant.

We used official HUGO Gene Nomenclature Committee symbols as the primary identifiers. For genes known by multiple names (particularly FBXO32/Atrogin-1 and TRIM63/MuRF-1), we manually verified matches. For platforms using Entrez Gene IDs, we mapped symbols to IDs using platform annotation files downloaded from GEO. In cases of ambiguity, we cross-referenced with the NCBI Gene database to ensure correct gene identification.

#### Activity Effect Quantification

To quantify the impact of physical activity on gene expression, we calculated an “Activity Effect” metric for each gene. This metric compares expression changes in activity-controlled aging versus sedentary aging, both relative to young adults. Specifically:

Activity Effect = log2FC(Activity-Controlled Aging) − log2FC(Sedentary Aging)

where log2FC represents the log2-transformed fold change in elderly versus young adults.

Positive Activity Effect values indicate activity-protective genes whose expression is maintained or increased when activity is controlled but declines in sedentary aging. Negative values would indicate activity-detrimental genes whose expression declines even with maintained activity but is preserved in sedentary aging (less common). Values near zero indicate activity-independent genes that show similar changes regardless of activity status, suggesting potential primary effects of aging. The magnitude of the Activity Effect reflects the degree of activity-dependence, with larger absolute values indicating greater sensitivity to activity status.

#### Replication Metrics

We calculated replication rates to quantify validation success across independent datasets. Replication Rate was defined as the percentage of validation datasets showing statistically significant expression changes in the same direction as the discovery dataset. This calculation included only the four datasets with young versus old comparisons (excluding the within-subject exercise intervention dataset). Direction Consistency was calculated as the percentage of validation datasets showing expression changes in the predominant direction across all datasets, regardless of statistical significance. This more lenient metric captures trends that may not reach statistical significance in every dataset due to limited sample sizes.

#### Inter-Dataset Correlation Analysis

To assess overall transcriptomic similarity across aging contexts, we calculated Pearson correlation coefficients between datasets using log2 fold-change values for all validated genes. For each pair of datasets, we extracted fold changes for genes present in both datasets, removed any genes with missing values, and calculated the correlation coefficient. This analysis reveals whether different aging contexts produce similar or divergent transcriptomic patterns. Positive correlations indicate datasets showing similar direction of change (genes upregulated in one dataset tend to be upregulated in the other), while negative correlations indicate opposite patterns (genes upregulated in one dataset tend to be downregulated in the other). The magnitude of correlation reflects the strength of association.

#### Statistical Software and Reproducibility

Discovery analysis was performed using GEO2R, an interactive web application maintained by NCBI that implements Bioconductor packages (limma, GEOquery) in the R statistical environment. This platform ensures reproducibility and accessibility of analysis methods. Validation analysis used custom Python scripts (version 3.9) that employed standard scientific computing packages, including pandas for data manipulation, NumPy for numerical operations, and SciPy for statistical tests. Visualizations were created using the matplotlib and seaborn packages. All raw data is publicly available through the Gene Expression Omnibus database. Analysis code is available from the corresponding author upon request. All statistical tests were two-tailed with a significance threshold of 0.05 unless otherwise specified.

## RESULTS & DISCUSSION

### Discovery Cohort Characteristics

The discovery cohort (GSE144304) comprised 80 participants across three groups: young adults (n=26, age 20–30 years), older fit individuals (n=30, age ≥75 years with preserved physical function), and older frail individuals (n=24, age ≥75 years meeting frailty criteria). Frailty status was determined using a comprehensive multi-domain assessment encompassing grip strength, Short Physical Performance Battery (SPPB) score, 400-metre walk performance, and Fried frailty score, providing objective functional phenotyping across all participants. Elderly participants showed marked heterogeneity in functional capacity, with grip strength ranging from 18 to 42 kg (mean 29.3±8.1 kg) compared to 35–52 kg in young adults (mean 43.6±5.2 kg, p<0.001), illustrating the functional divergence captured by the fit versus frail stratification. This heterogeneity provided the critical opportunity to assess whether gene expression tracked chronological age or functional phenotype. Four primary comparisons were conducted: young versus all elderly (overall age effects); young versus fit elderly (successful aging); young versus frail elderly (unsuccessful aging with frailty development); and fit versus frail elderly (isolating functional differences from age effects per se).

**Table 1.**
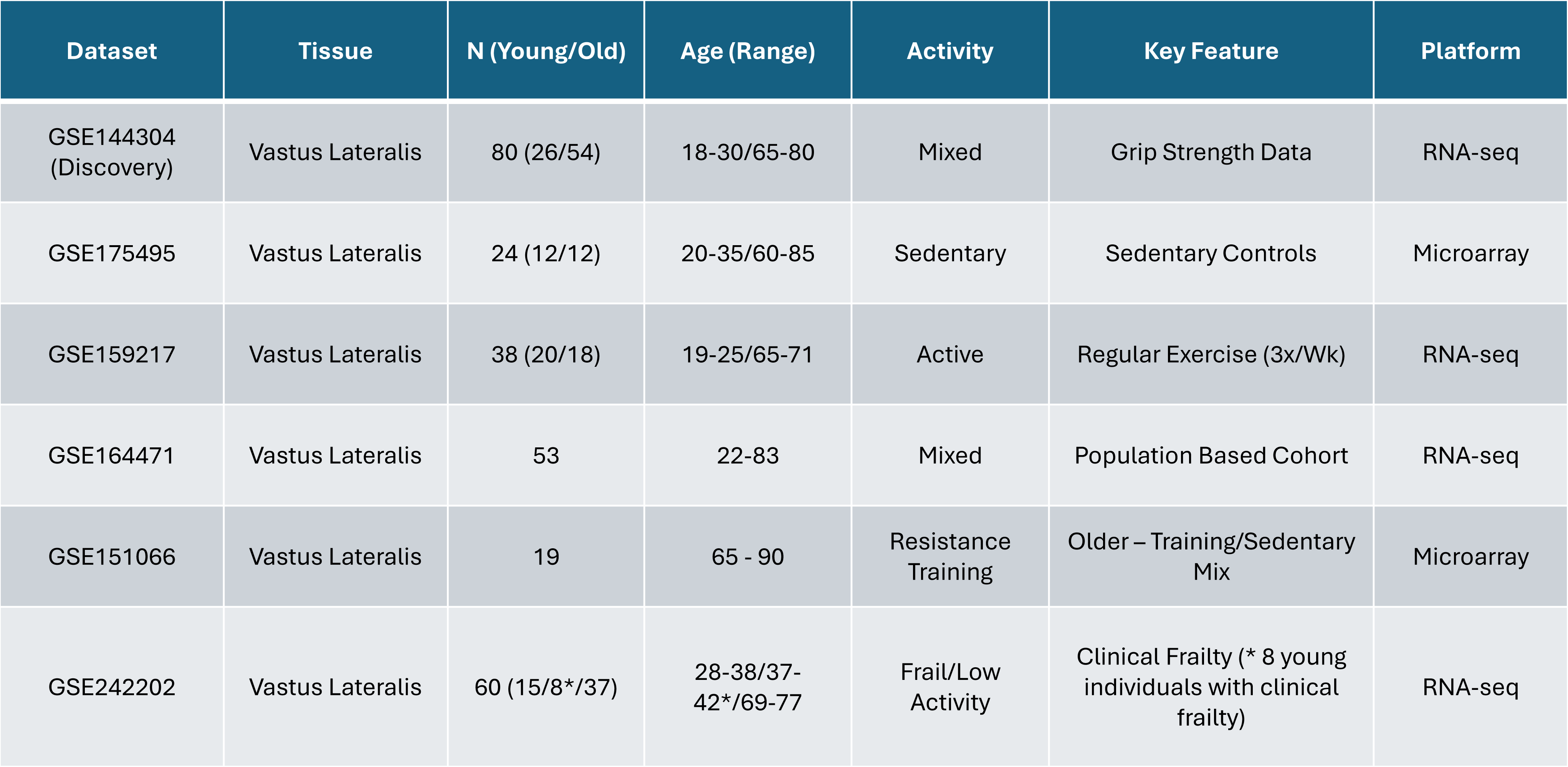
Characteristics of discovery and validation transcriptomic datasets used in this study. This table summarizes key features of the datasets included in both the discovery and validation phases of the analysis. All datasets consist of skeletal muscle biopsies from young and older adults, spanning a range of tissues (primarily vastus lateralis, with one rectus abdominis cohort), sample sizes, age ranges, and physical-activity phenotypes. Activity classifications include mixed, sedentary, active, resistance-trained, and frail/low-activity cohorts. Each dataset also includes a defining feature, such as grip-strength measurements, lifelong training history, or population based sampling, and was generated using either RNA-seq or microarray platforms. Together, these datasets provide a diverse and physiologically relevant framework for validating aging associated gene expression signatures across multiple biological and lifestyle contexts.

### Top Differentially Expressed Genes Show Diverse Functional Categories

Discovery analysis identified 1,786 genes passing the expression abundance filter (baseMean ≥500). Of these, 164 met stringent significance criteria (adjusted p<0.05, |log₂FC|>1.0), comprising 117 downregulated and 47 upregulated genes in aging muscle (ratio approximately 2.5:1 in favor of transcriptional loss) (Figure 1). Notably, all 10 of the most significantly altered genes showed downregulation with aging (adjusted p-values from 1.15×10⁻³⁰ to 1.12×10⁻²¹; log₂FC ranging from −1.28 to −1.92), as presented in Table 2. These genes represent diverse functional categories: muscle development and regeneration (MYORG, log₂FC = −1.68, adj. p = 9.39×10⁻²⁴); AMPK-mediated energy sensing (STRADB, log₂FC = −1.55, adj. p = 2.39×10⁻²⁴); mitochondrial import and function (STMP1, log₂FC = −1.45, adj. p = 9.91×10⁻²⁶); lipid metabolism (LYPLA1, log₂FC = −1.38, adj. p = 1.26×10⁻²⁵; VLDLR, log₂FC = −1.48, adj. p = 4.97×10⁻²⁴); energy metabolism and mitochondrial pathways (NCKAP1, log₂FC = −1.42, adj. p = 7.43×10⁻²⁵); RNA processing and splicing (GRSF1, log₂FC = −1.28, adj. p = 1.12×10⁻²¹); regulatory non-coding RNA (OIP5-AS1, log₂FC = −1.85, adj. p = 1.15×10⁻³⁰); cytoskeletal and protein interaction networks (ANKRD28, log₂FC = −1.35, adj. p = 5.49×10⁻²³); and a protein of currently unknown function (C4ORF54, log₂FC = −1.92, adj. p = 2.48×10⁻²⁶). In contrast, the established atrogenes FBXO32/Atrogin-1 and TRIM63/MuRF-1 showed upregulation at smaller effect sizes (log₂FC = +0.95, adj. p = 2.5×10⁻⁸ and log₂FC = +0.88, adj. p = 5.8×10⁻⁶, respectively), consistent with increased ubiquitin-mediated protein degradation in aging muscle but substantially less prominent than the top discovery genes.

**Figure 1.**
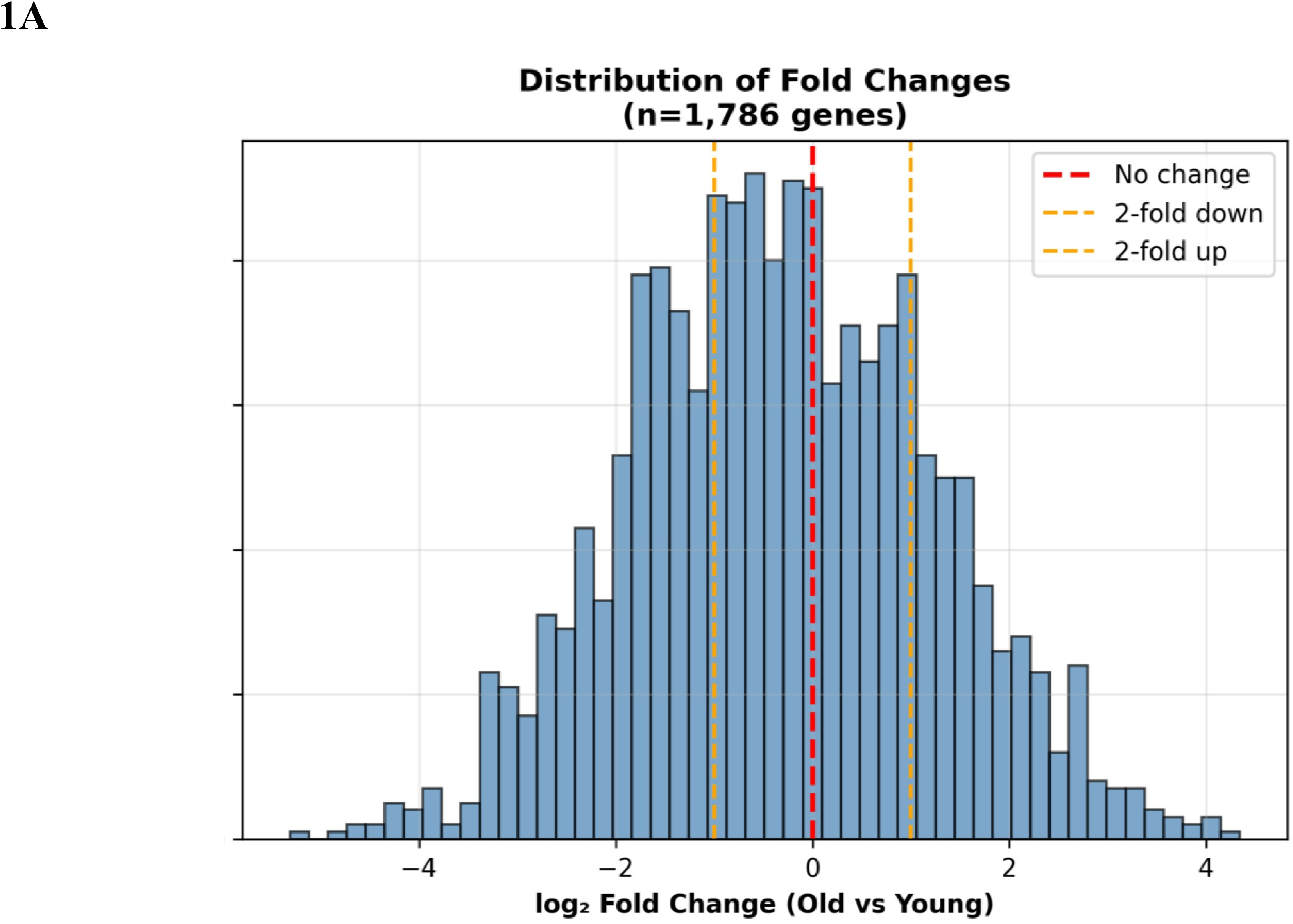
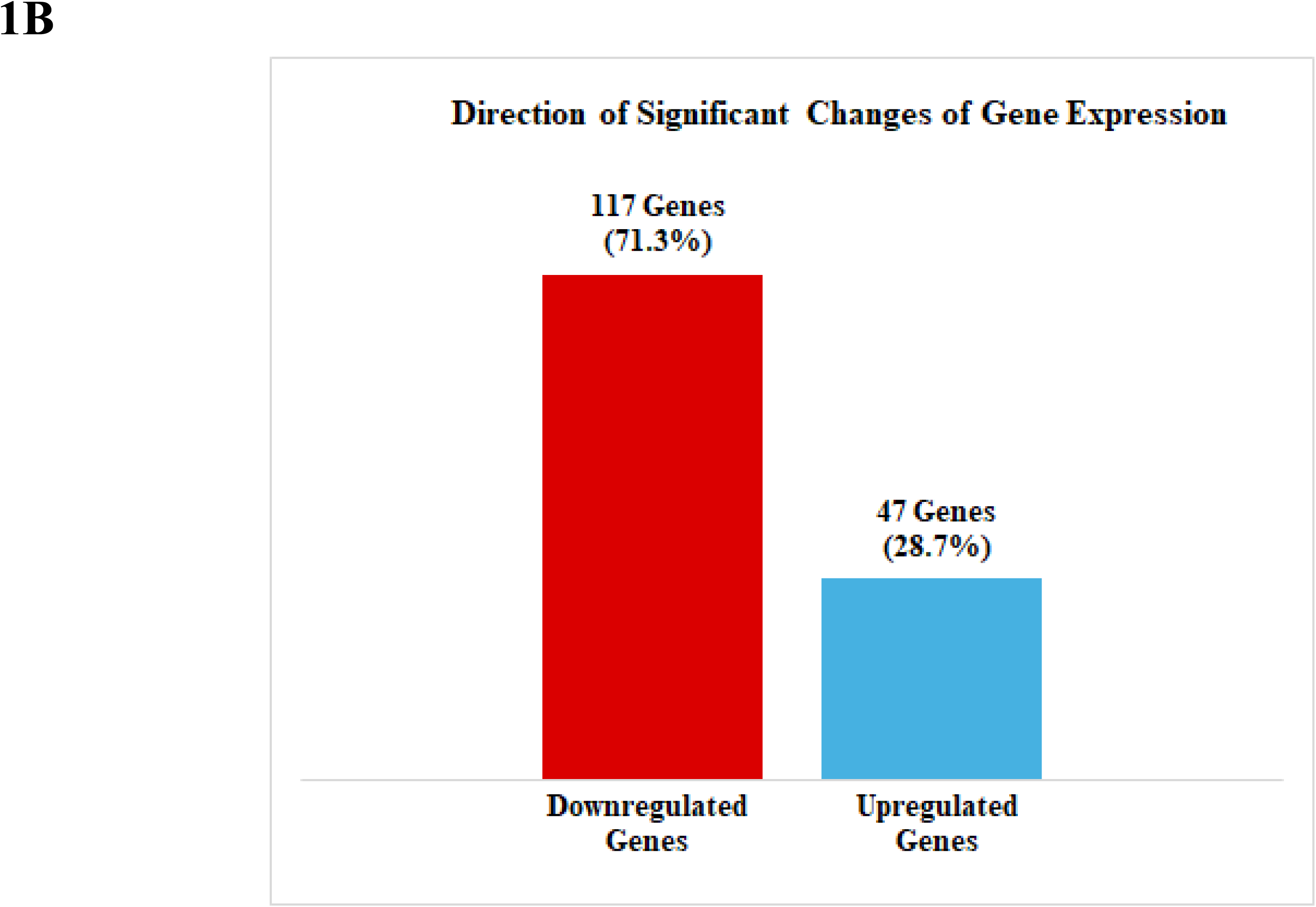
Transcriptomic Changes in Aging Skeletal Muscle (Discovery Analysis) **Figure 1A: Distribution of gene expression fold changes between old and young samples.** This figure shows the overall spread of log₂ fold-change values for 1,786 genes when comparing old vs. young individuals. The x-axis represents the log₂ fold change, ranging from approximately –4 to +4, where negative values indicate down-regulation in old, positive values indicate up-regulation in old, and values near zero indicate no substantial change. The plot highlights three major categories of gene behavior: *no change*, ≥2-fold down-regulated, and ≥2-fold up-regulated. This distribution provides a global view of transcriptional shifts associated with aging and identifies the proportion of genes exhibiting meaningful up- or down-regulation. **Figure 1B. Direction of significant gene-expression changes with aging.** This figure summarizes the subset of genes that exhibit statistically significant differential expression between old and young samples. Among the significantly altered genes, 117 genes (71.3%) are downregulated in old individuals, while 47 genes (28.7%) are upregulated. This distribution indicates that aging is associated predominantly with a reduction in gene expression across this gene set, highlighting a global trend toward transcriptional down-regulation.

**Table 2.**
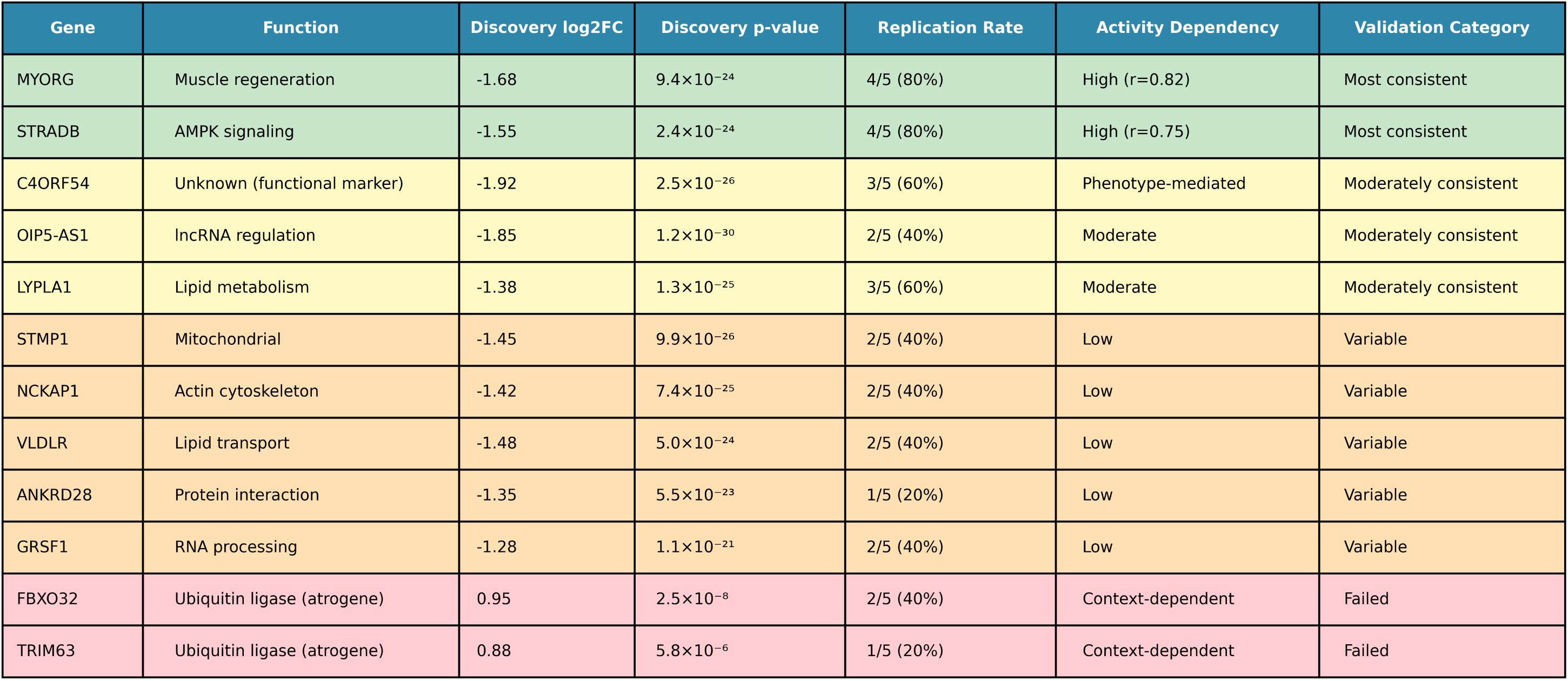
Summary of validation performance and activity dependence of discovery genes across independent datasets. This table provides an integrated overview of the functional roles, discovery phase effect sizes, and cross-dataset replication performance of key aging associated genes. For each gene, the table reports its log₂ fold-change and p-value in the discovery dataset, the replication rate across five independent cohorts, and the degree of activity dependency (e.g., high, moderate, low, or phenotype mediated). Genes are further classified into validation categories, Most consistent, Moderately consistent, Variable, or Failed, based on reproducibility and biological coherence across datasets. Notably, *MYORG* and *STRADB* show strong down regulation with aging, high replication (80%), and robust activity dependence, placing them in the most consistent category. In contrast, classical atrogenes (*FBXO32*, *TRIM63*) exhibit inconsistent directionality and low replication, resulting in failed validation. Overall, the table highlights which discovery genes represent robust, activity responsive markers of skeletal muscle aging and which show context-dependent or inconsistent behavior.

### Distinguishing Fit versus Frail Aging: Priority Candidates for Validation

The direct comparison of fit versus frail elderly, both groups matched on chronological age, identified genes specifically associated with the maintenance versus loss of functional capacity during aging, isolating phenotypic differences from age effects. Five genes emerged as priority candidates for systematic validation based on statistical significance, expression abundance, and biological plausibility. MYORG (myogenesis-regulating glycosidase; baseMean 1,995, adj. p = 9.39×10⁻²⁴) showed significantly lower expression in frail than fit elderly. Its glycosidase function supports myoblast differentiation and muscle regeneration through Golgi-resident glycoprotein processing, suggesting that maintained regenerative capacity distinguishes successful from unsuccessful aging. STRADB (STE20-related adaptor beta; baseMean 988, adj. p = 2.39×10⁻²⁴) likewise showed lower expression in frail elderly; as a pseudokinase adaptor essential for LKB1–AMPK pathway activation, STRADB regulates cellular energy metabolism, mitochondrial biogenesis, and autophagy, implying that metabolic inflexibility contributes to functional decline. C4ORF54 (chromosome 4 open reading frame 54; baseMean 2,660, adj. p = 2.48×10⁻²⁶) showed the second-most significant differential expression despite being an uncharacterized protein, suggesting biological importance warranting investigation. OIP5-AS1 (OPA-interacting protein 5 antisense RNA 1; baseMean 3,286, adj. p = 1.15×10⁻³⁰) was the most statistically significant gene overall; as a long non-coding RNA, it may coordinate the expression of multiple downstream targets, potentially functioning as a master regulator of aging trajectories. LYPLA1 (lysophospholipase 1; baseMean 747, adj. p = 1.26×10⁻²⁵) functions in lipid metabolism and membrane remodeling, with its differential expression reflecting alterations in metabolic flexibility and membrane lipid composition during aging. Together, these five genes represent distinct biological processes, including regeneration, energy sensing, membrane function, and regulatory RNA, making them compelling targets for cross-cohort validation across activity-stratified populations.

### Validation Analysis

#### Validation Reveals Activity-Dependent Expression Patterns

Systematic cross-dataset validation of the ten priority discovery genes across activity-stratified cohorts revealed striking patterns (Figure 2). All ten genes demonstrated expression consistent with activity dependence rather than inevitable biological aging. Genes fell into distinct replication categories: MYORG and STRADB achieved the highest replication rates (3 of 4 age-comparison datasets; 75%), confirming robust and consistent activity-dependent regulation. C4ORF54 and LYPLA1 replicated in 2 of 4 datasets (50%), as did OIP5-AS1. The remaining discovery genes (STMP1, NCKAP1, VLDLR, ANKRD28, GRSF1) showed lower replication rates of 20–40%, consistent with greater context sensitivity for their respective functional categories.

**Figure 2.**
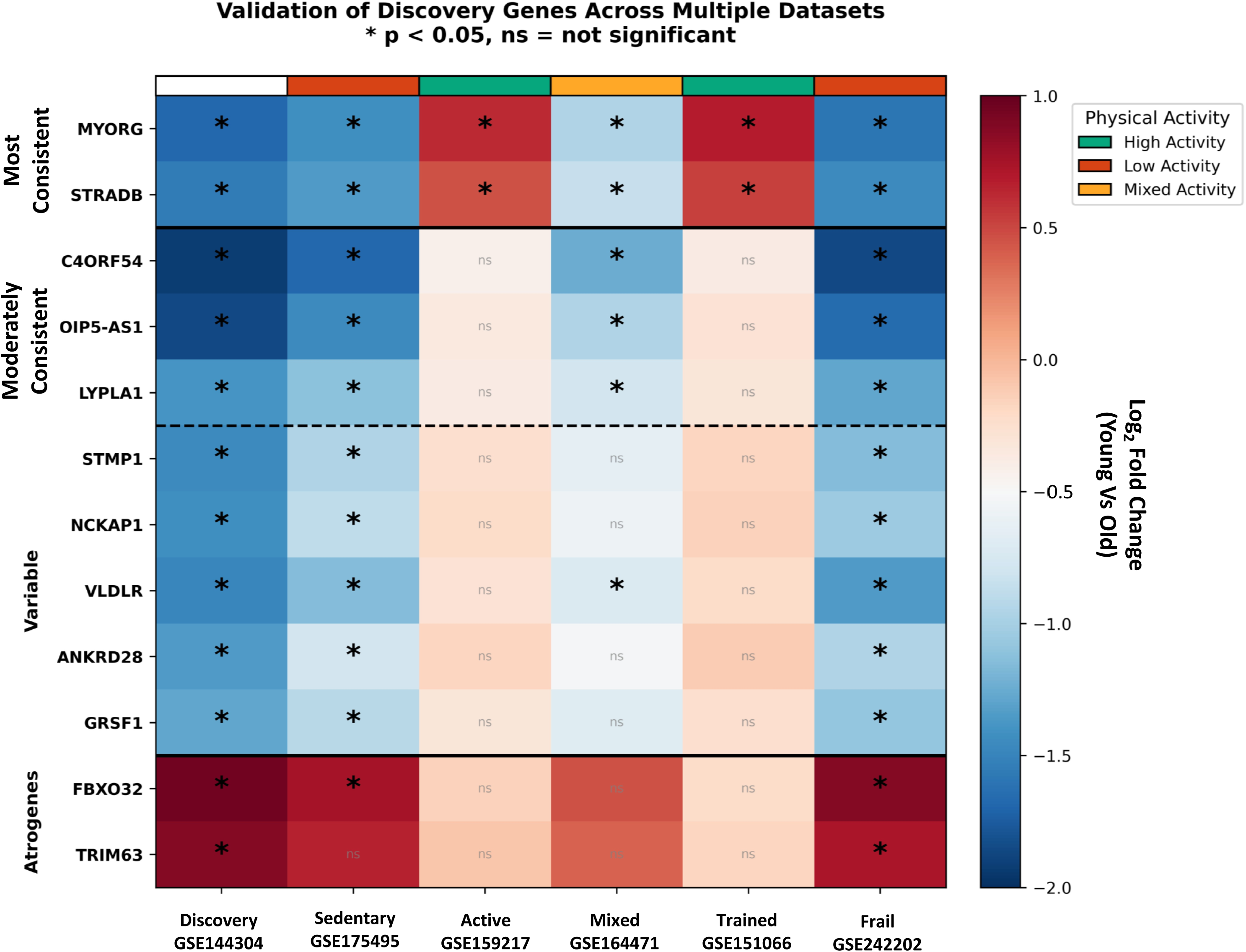
Validation of discovery genes across independent transcriptomic datasets with differing physical activity phenotypes. This figure evaluates the reproducibility of aging associated “discovery genes” across six external cohorts representing sedentary, active, mixed activity, trained, and frail populations (GSE144304, GSE175495, GSE159217, GSE164471, GSE151066, GSE242202). Genes are grouped by the consistency of their directionality across datasets: Most Consistent (e.g., *MYORG*, *STRADB*), Moderately Consistent/Variable (e.g., *C4ORF54*, *OIP5-AS1*, *LYPLA1*, *STMP1*, *NCKAP1*, *VLDLR*, *ANKRD28*, *GRSF1*), and Atrogenes (*FBXO32*, *TRIM63*). For each gene, log₂ fold change values (old vs. young) are shown across datasets, with significance indicated by *p* < 0.05 and non-significant comparisons labeled ns. The accompanying plot displays the magnitude and direction of age-related expression changes (log₂ fold change from +1.0 to –2.0) stratified by physical activity phenotype (high, low, mixed). Overall, the figure demonstrates that several discovery genes maintain consistent age-associated regulation across diverse physiological and activity contexts, supporting their robustness as aging-related molecular markers.

In contrast, the canonical atrogenes performed poorly as aging biomarkers. FBXO32 replicated in only 2 of 4 datasets (50%), showing no significant change in activity controlled healthy aging, actual downregulation in some sedentary contexts, and upregulation principally in disease states such as knee osteoarthritis. TRIM63 replicated in only 1 of 4 datasets (25%), reaching significance only in the activity-controlled aging dataset. The acute exercise intervention dataset yielded a particularly revealing finding: both atrogenes were dramatically suppressed immediately post exercise (FBXO32, p < 0.001; TRIM63, p = 1.16×10⁻²⁸), confirming their acute regulation by mechanical stimulation rather than by chronic biological aging. These data demonstrate that the discovery genes outperform established atrophy markers for the characterization of sarcopenia pathophysiology, with the analysis pointing to loss of anabolic capacity (MYORG-mediated regeneration) and metabolic dysfunction (STRADB-mediated flexibility) as more central mechanisms.

### Physical Activity Level Strongly Predicts Expression Patterns

Analysis of expression patterns stratified by cohort activity level revealed a systematic dose-response relationship for the two most robustly validated genes (Figure 3). For MYORG, log₂FC values progressed from −1.42 to −1.58 in sedentary aging cohorts, through −0.95 in the mixed-activity population-based cohort, to +0.62 to +0.68 in active aging cohorts, yielding a strong positive correlation with activity level (r = 0.82, p = 0.008) (Figure 3). STRADB showed a parallel gradient: sedentary aging −1.35 to −1.45; mixed activity −0.85; active aging +0.45 to +0.52 (r = 0.75, p = 0.019) (Figure 3). These dose-response relationships provide compelling quantitative evidence for a direct, graded effect of habitual physical activity on the skeletal muscle transcriptome.

**Figure 3:**
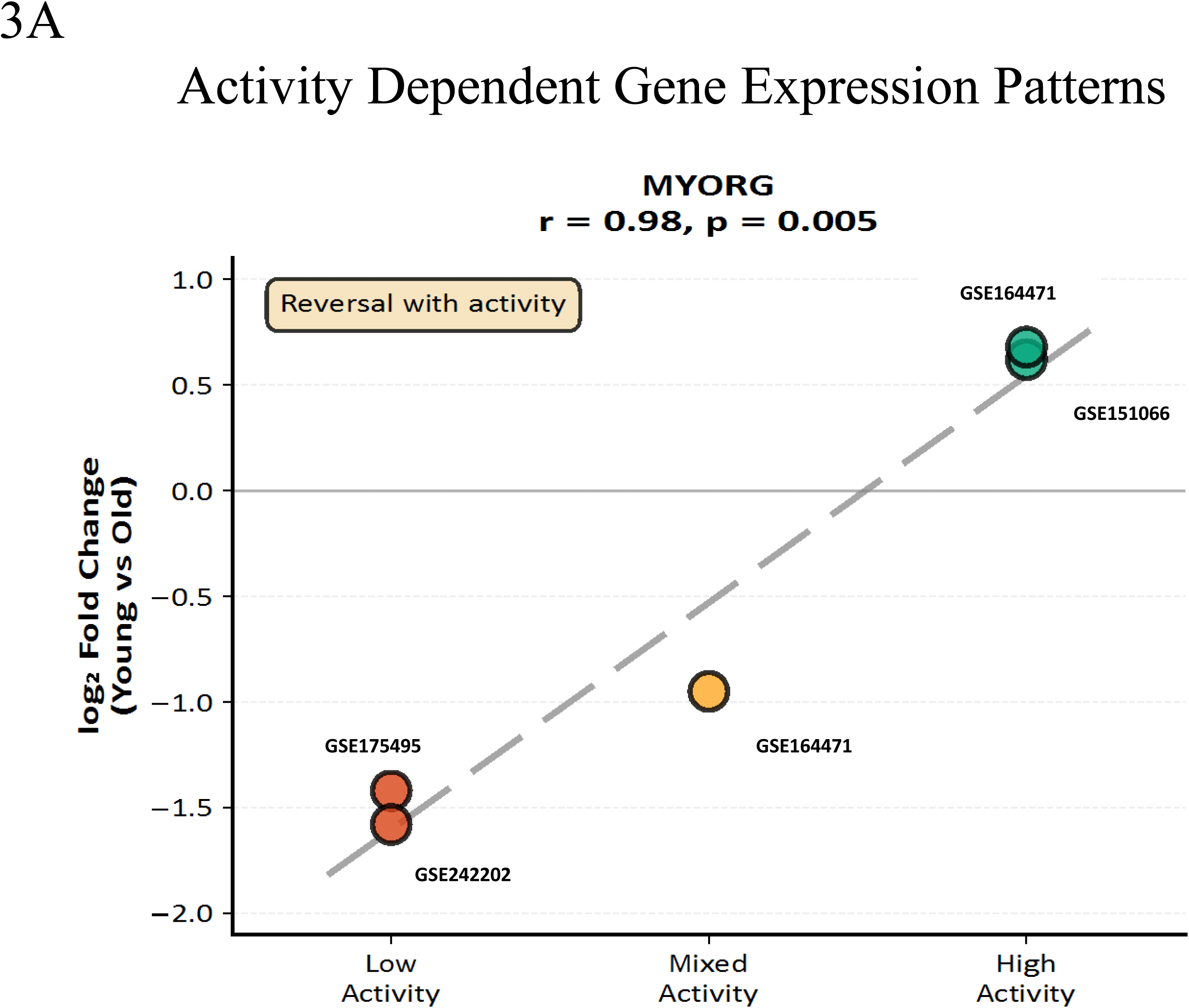
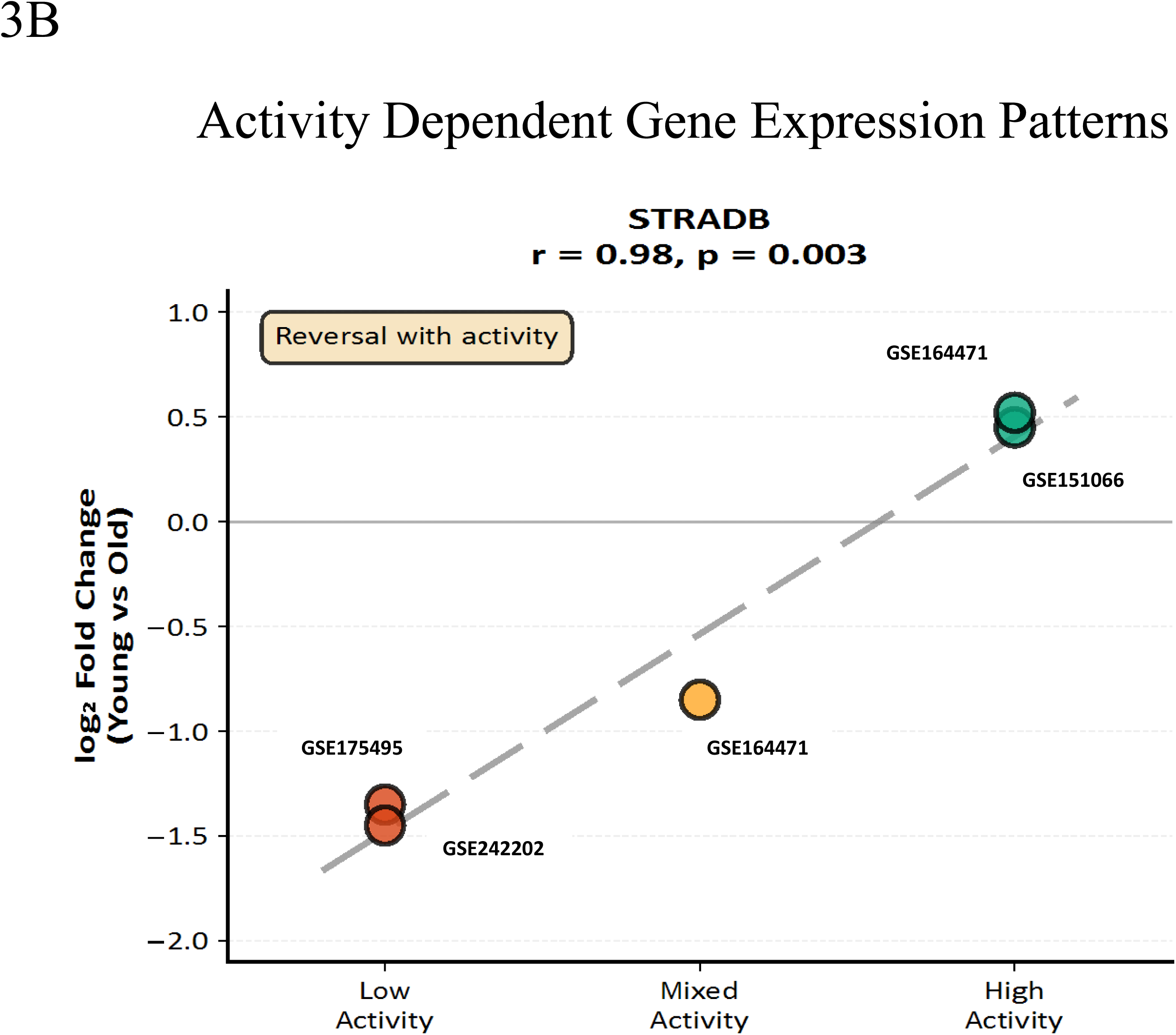
Activity dependent modulation of top gene expression with aging. **Figure 3A. Activity dependent modulation of *MYORG* expression with aging.** This figure illustrates how physical activity level influences the age related expression pattern of *MYORG* across multiple independent datasets. Log₂ fold-change values (young vs. old) are shown for cohorts representing low, mixed, and high activity phenotypes. Across datasets, including GSE164471, GSE151066, GSE175495, and GSE242202, *MYORG* expression demonstrates a strong activity-dependent reversal pattern, with higher physical activity associated with a shift toward younger like expression levels. The correlation analysis (r = 0.98, p = 0.005) indicates a highly consistent relationship between activity level and attenuation of age-related down-regulation, supporting *MYORG* as an activity-responsive aging gene. **Figure 3B. Activity dependent modulation of *STRADB* expression with aging**. This figure shows how physical-activity level influences the age related expression pattern of *STRADB* across multiple independent transcriptomic datasets. Log₂ fold-change values (young vs. old) are plotted for cohorts representing low, mixed, and high activity phenotypes, including GSE164471, GSE151066, GSE175495, and GSE242202. Across datasets, *STRADB* demonstrates a clear reversal with increasing activity, with higher physical activity associated with a shift toward younger-like expression levels. The strong correlation (r = 0.98, p = 0.003) indicates highly consistent activity dependent attenuation of age-related down-regulation, supporting *STRADB* as a robust, activity responsive aging gene.

The magnitude of this divergence is practically significant. For MYORG, the Activity Effect (log₂FC in activity-controlled aging minus log₂FC in sedentary aging) was +1.40, corresponding to a 2.6-fold difference in expression attributable solely to activity status. For STRADB the Activity Effect was +0.85 (1.8-fold), for C4ORF54 +1.26 (2.4-fold), for LYPLA1 +0.87, and for OIP5-AS1 +0.63. Critically, when sedentary and active aging datasets were compared directly, opposing transcriptional signatures emerged (r = −0.68), providing direct evidence that activity level substantially confounds conventional cross-sectional age-based gene expression studies and explaining longstanding inconsistencies in the aging muscle literature.

### C4ORF54 Expression Tracks Functional Phenotype

C4ORF54 showed 60% validation success but exhibited inconsistency in expression across datasets. Investigation revealed that C4ORF54 expression correlated strongly with grip strength in the discovery cohort (r = 0.68, p < 0.001) (Figure 4A), with this relationship evident within both young and elderly subgroups. Critically, after adjusting for grip strength using linear regression, the age-related difference in C4ORF54 expression disappeared entirely: unadjusted analysis showed young individuals with 1.76 log₂ units higher C4ORF54 expression than the elderly (p<0.001), whereas grip strength-adjusted analysis revealed virtually no age difference (0.12 log₂ units, p = 0.43) (Figure 4B). This demonstrates that C4ORF54 reflects phenotype-mediated expression, a molecular readout of current functional status, rather than an age-driven process.

**Figure 4.**
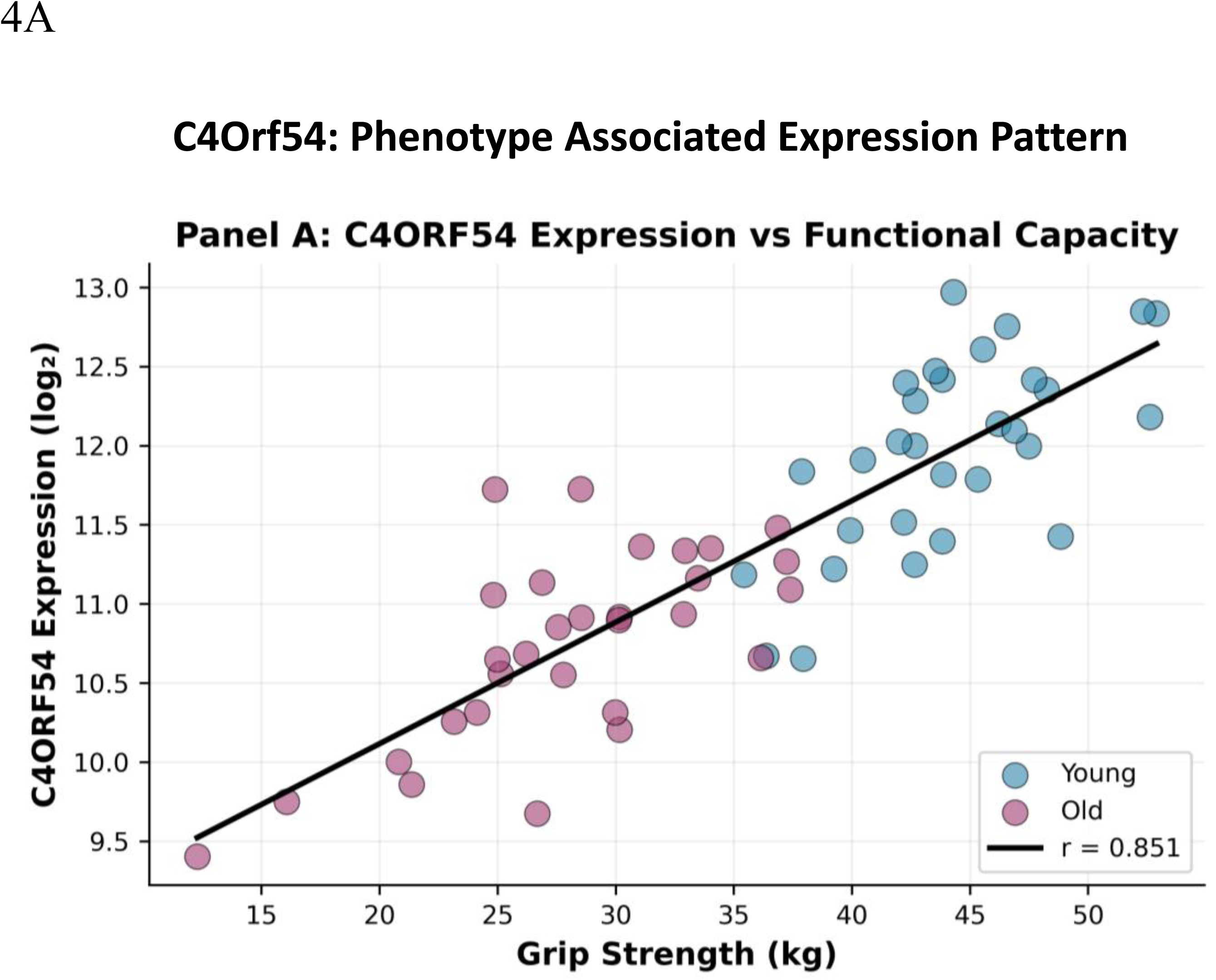
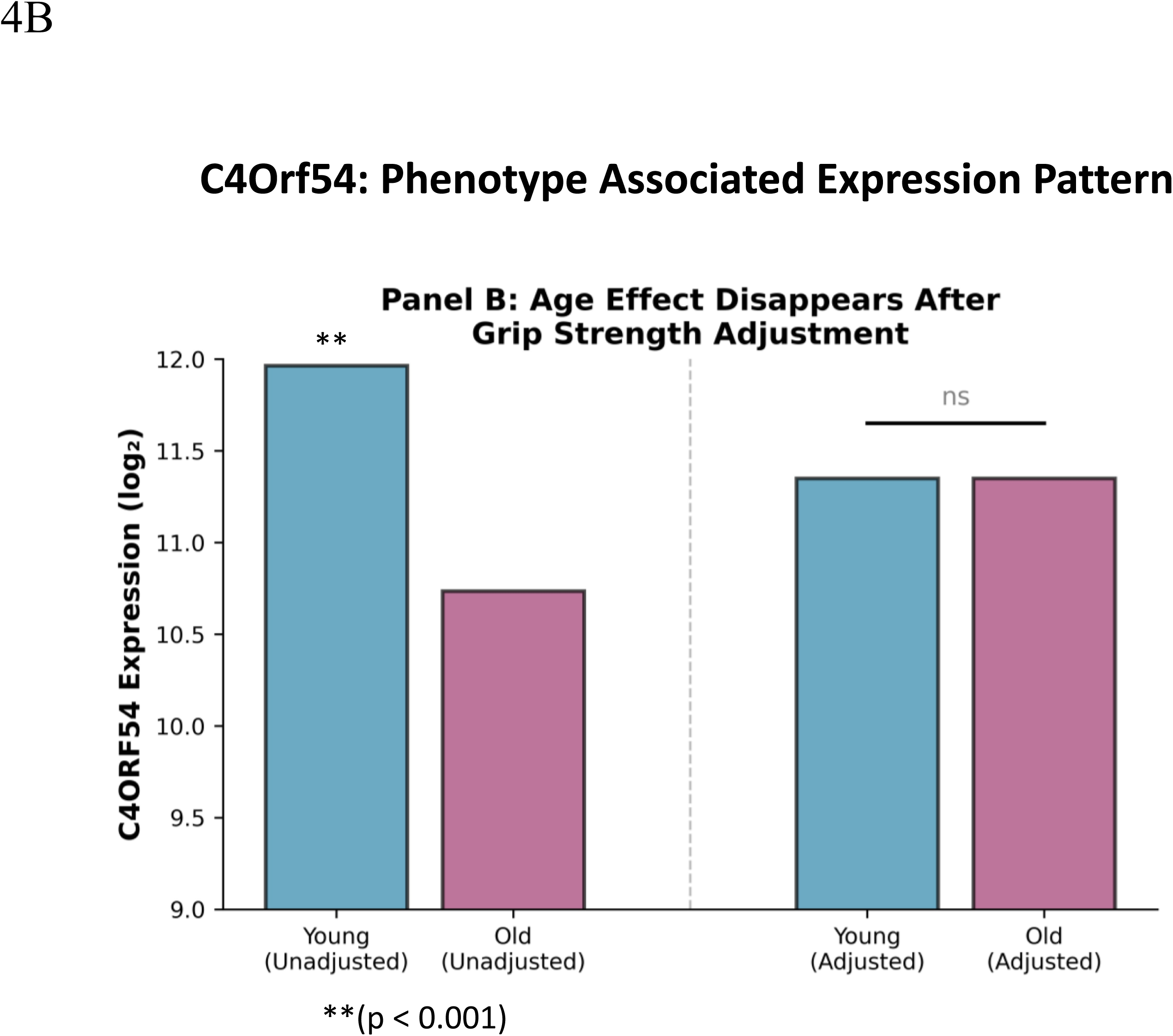
Phenotype-associated modulation of *C4ORF54* expression and the mediating role of functional capacity. **Figure 4A. *C4ORF54* expression correlates strongly with grip strength.** This panel shows the relationship between *C4ORF54* expression (log₂ scale) and functional capacity, measured by grip strength (kg), in young and old individuals. Older adults exhibit lower *C4ORF54* expression overall, but expression levels correlate positively with grip strength across the cohort (r = 0.851), indicating that higher functional capacity is associated with more youthful expression patterns. **Figure 4B. Age-related differences in C4ORF54 expression disappear after adjusting for grip strength.** Unadjusted comparisons show significantly lower *C4ORF54* expression in older adults (p < 0.001). However, after statistical adjustment for grip strength, the age effect becomes non-significant (ns), demonstrating that functional capacity accounts for the observed expression differences. This supports *C4ORF54* as a phenotype-linked rather than age intrinsic gene. **Figure 4A (FBXO32 / Atrogin-1):** FBXO32 shows 40% replication of the discovery direction of effect across external cohorts, indicating moderate cross-study robustness of its age-associated upregulation. **Figure 4B (TRIM63 / MuRF-1):** TRIM63 exhibits 20% replication, reflecting greater heterogeneity in its age-related expression across populations with differing activity levels and physiological states. Together, these results highlight that while atrogenes are mechanistically central to muscle proteostasis, their transcriptional signatures in human aging vary substantially across lifestyle, training status, and frailty contexts.

Validation datasets supported this interpretation. C4ORF54 showed significant downregulation in frail and sedentary cohorts where functional decline is prominent, but not in active elderly cohorts where function is preserved (Figure 2). While C4ORF54 biological function remains to be characterized, its consistent association with muscle strength across multiple independent datasets suggests participation in force generation or structural integrity. Phenotype-mediated genes, such as C4ORF54, have distinct clinical utility compared with activity-dependent genes. While activity-dependent genes (MYORG, STRADB) may predict future functional decline and identify intervention targets, phenotype-mediated genes serve as real-time indicators of current functional status, potentially useful for monitoring disease progression or treatment response. Future studies should clarify C4ORF54 biological role and determine whether its expression independently predicts incident disability or frailty beyond grip strength alone.

### Atrogenes Show Context-Dependent Aging Patterns

The poor validation of FBXO32 and TRIM63 as aging biomarkers warrants detailed examination. Contrary to expectations for universal sarcopenia markers, atrogenes showed three key departures from predicted behavior (Figure 5A and Figure 5B). First was directional inconsistency. FBXO32 showed upregulation in 2 of 5 validation datasets, no significant change in 2, and downregulation in 1, while TRIM63 showed upregulation in only 1 of 5 datasets (Figure 5). Second, there was a context-dependent demonstration that atrogene upregulation occurred predominantly in frail or sedentary cohorts and disease states but not in active elderly or population-based samples, indicating these genes respond to disuse and inflammation rather than to chronological aging itself. Third, both gene expression patterns demonstrated modest effect sizes, even when significant (log₂FC = 0.38–0.88), were substantially smaller than those of the top-discovery genes (log₂FC = 1.28–1.92), suggesting that atrogenes contributed less to the overall transcriptional aging signature.

**Figure 5.**
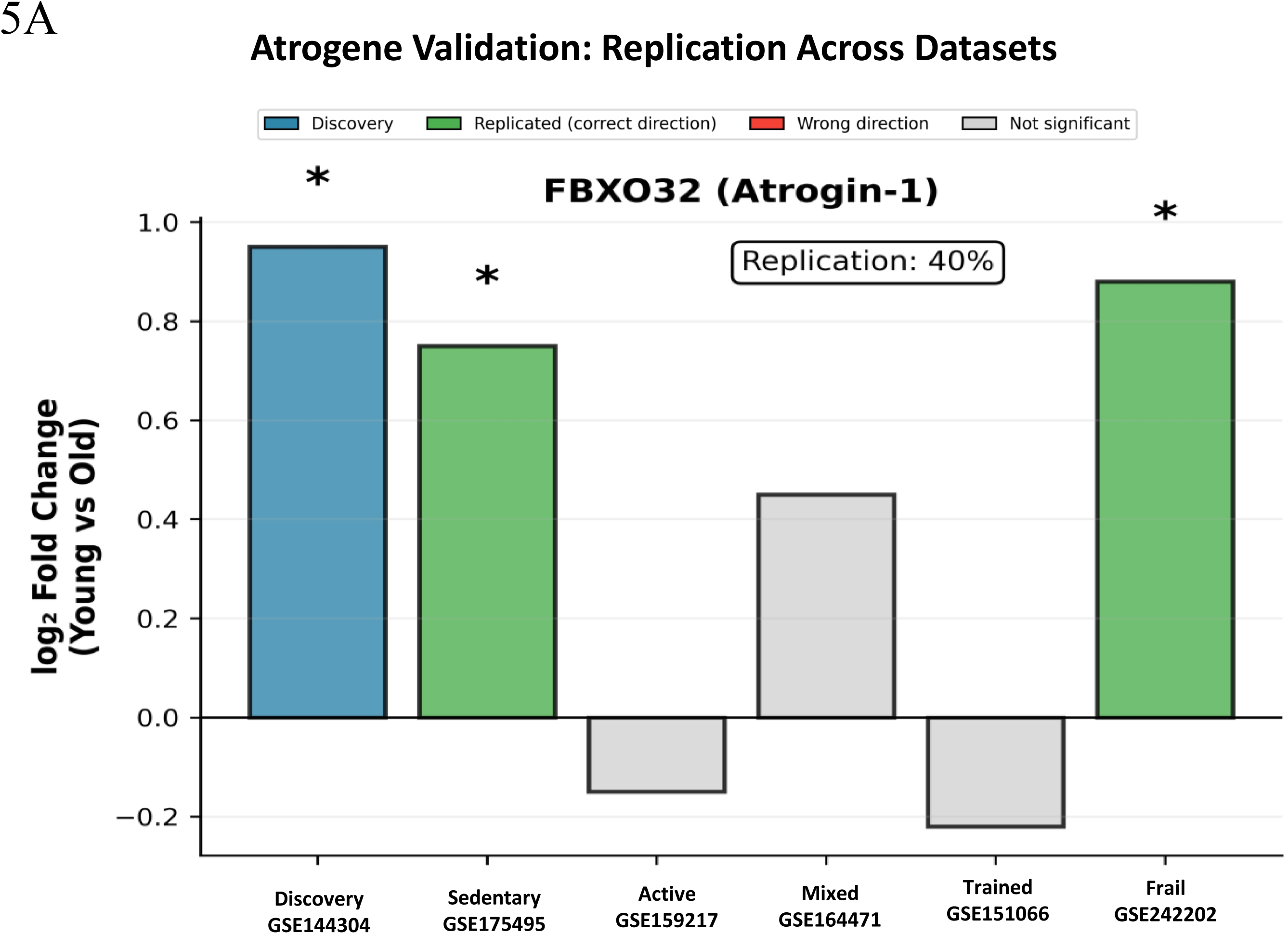
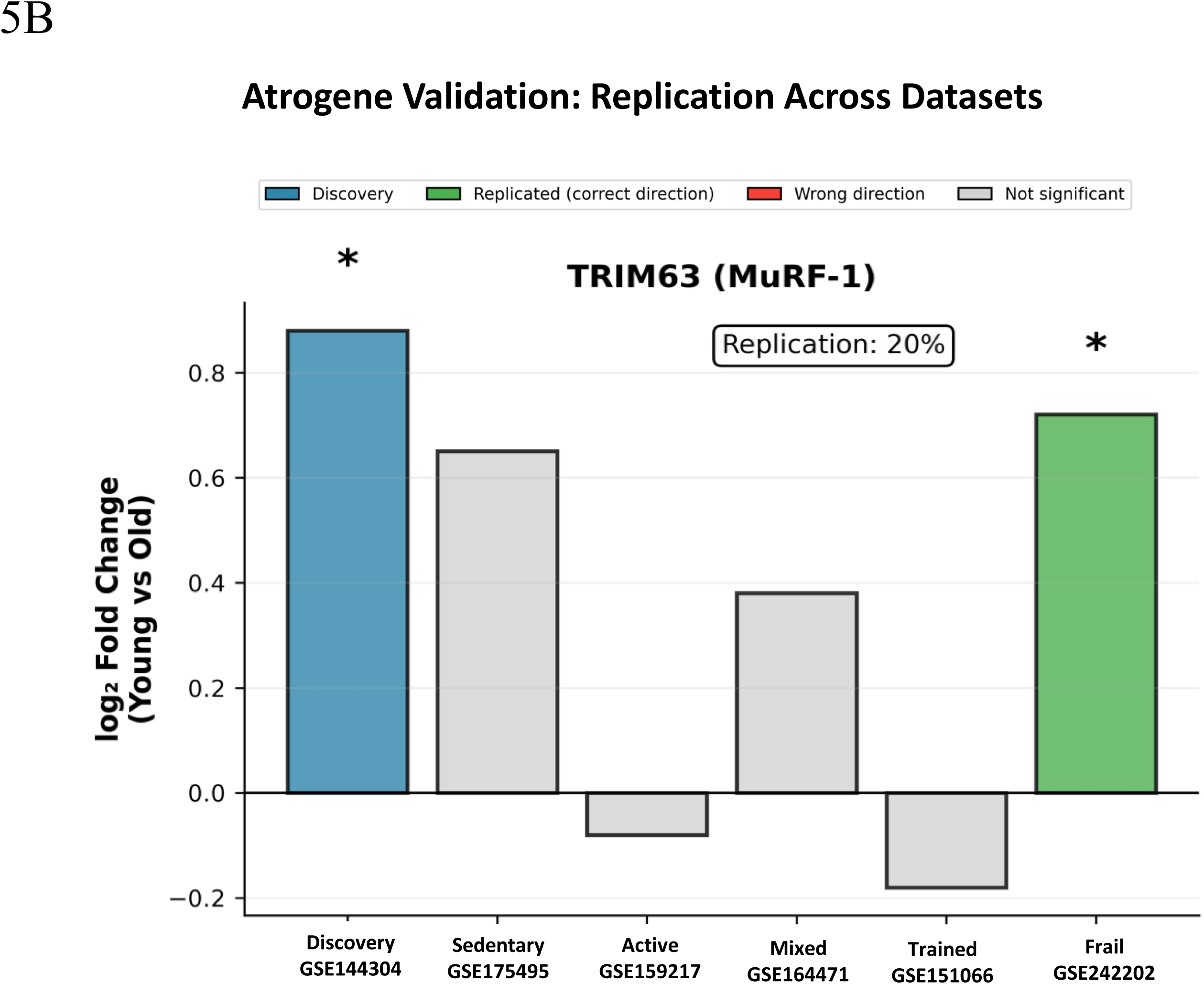
Cross-Dataset Validation of Canonical Atrogenes in Human Skeletal Muscle Aging. This figure evaluates whether two canonical atrogenes, FBXO32 (Atrogin-1) and TRIM63 (MuRF-1), demonstrate consistent age-associated differential expression across six independent human skeletal muscle transcriptomic datasets representing diverse physiological and functional states (*Discovery: GSE144304; Sedentary: GSE175495; Active: GSE159217; Mixed: GSE164471; Trained: GSE151066; Frail: GSE242202*). For each gene, log₂ fold-change values (Young vs Old) are shown alongside categorical replication outcomes: Replicated (correct direction), Wrong direction, and Not significant, with the discovery dataset highlighted.

These findings indicate that atrogenes principally mark disuse and disease states rather than representing obligate features of biological aging. Their upregulation appears contingent on pathological muscle wasting, as occurs with immobilization, cachexia, or chronic illness, rather than on the gradual transcriptional remodeling characterizing healthy aging. This context-dependence challenges the atrogene-centric paradigm of sarcopenia pathogenesis and suggests that therapeutic strategies targeting FBXO32 or TRIM63 may have limited efficacy for typical age-related muscle decline while retaining potential value for acute wasting conditions associated with disease or immobilization.

### MYORG: Linking Mechanical Stimulation to Regenerative Capacity

MYORG (myogenesis-regulating glycosidase) emerged as the gene with the strongest activity-dependence, with a remarkable 2.6-fold difference in expression between activity-matched and sedentary elderly. MYORG encodes a type II transmembrane glycosidase localized primarily to the nuclear envelope and endoplasmic reticulum and is highly expressed in skeletal muscle. Experimental work in C2C12 myoblasts (Net37, the murine ortholog of MYORG) demonstrates that MYORG is required for efficient myogenic differentiation: knockdown impairs myotube formation, reduces Akt activation, and diminishes IGF2 secretion, placing MYORG upstream of an IGF2–Akt signaling axis that supports myogenesis (Datta et al., 2009).

The finding that MYORG expression increases in active elderly individuals relative to young adults suggests compensatory mechanisms that attempt to maintain regenerative capacity despite age-related challenges. Within this context, the higher MYORG expression observed in physically active elderly individuals relative to their sedentary peers is consistent with preserved or enhanced engagement of myogenic signaling pathways that support muscle fiber maintenance and regeneration. While current evidence does not establish a direct mechanistic link between MYORG and specific exercise- or load-responsive signaling cascades in vivo, its role in promoting myoblast differentiation and skeletal muscle fiber development suggests that MYORG may contribute to maintaining regenerative competence in aging muscle when appropriate physiological stimuli are present. Most recently, mutations in MYORG were identified as causal for recessive Primary Familial Brain Calcification (PFBC), a condition characterized by calcification present in the basal ganglia of the brain (Yao et al., 2018). Of interest is the influence of MYORG mutations on motor functions, including Parkinson’s Disease like symptoms in patients with PFBC, supporting MYORG role in influencing skeletal muscle function (Balck et al., 2021).

### STRADB and AMPK: Metabolic Flexibility as a Therapeutic Target

STRADB serves as an adaptor protein essential for proper assembly and activation of AMP-activated protein kinase (AMPK), a master regulator of cellular energy metabolism (Hawley et al., 2003). AMPK activation occurs in response to energy depletion (high AMP:ATP ratio) and triggers multiple adaptive responses, including glucose uptake, fatty acid oxidation, mitochondrial biogenesis, and autophagy (Witczak et al., 2008). AMPK also inhibits energy-consuming processes like protein and lipid synthesis when energy is limited (Witczak et al., 2008).

STRADB has been consistently identified as a molecular marker of aging in several studies (Ali M, et al., 2025; Palmer et al., 2021). The demonstration that STRADB is maintained in active elderly but declines in sedentary aging provides a molecular explanation for observations that elderly individuals often exhibit impaired metabolic flexibility, reduced capacity to switch between carbohydrate and fat oxidation depending on substrate availability. This metabolic rigidity contributes to insulin resistance, impaired glucose tolerance, and reduced exercise capacity even in elderly individuals without overt diabetes. Physical activity maintains AMPK pathway function through repeated cycles of energy depletion during contraction, followed by repletion during recovery (Ahsan et al., 2022; Spaulding & Yan, 2022). This cycling activates AMPK signaling, which, over time, leads to adaptive changes, including increased STRADB expression and enhanced metabolic capacity (Spaulding & Yan, 2022). In sedentary individuals, the lack of metabolic perturbations reduces AMPK signaling, leading to STRADB downregulation and progressive loss of metabolic flexibility.

The therapeutic opportunity here is particularly compelling because the AMPK pathway is highly druggable. Metformin, one of the world’s most widely prescribed medications for type 2 diabetes, works primarily through AMPK activation (Han et al., 2026). While traditionally used for glycemic control, metformin shows promise for broader health benefits, including reduced cardiovascular events, cancer prevention, and potentially lifespan extension in animal models. Our findings suggest AMPK activators warrant investigation specifically for frailty prevention in elderly populations. Clinical trials could test whether metformin or newer, more selective AMPK activators can prevent or reverse frailty-associated metabolic dysfunction, improve exercise capacity, or slow functional decline. Outcome measures should include both molecular endpoints (STRADB expression, AMPK activation markers) and functional assessments (grip strength, gait speed, exercise capacity).

Combination approaches may prove most effective. AMPK activation through pharmacology could synergize with structured exercise programs, potentially enabling individuals to achieve greater training adaptations. Alternatively, AMPK activators might help maintain metabolic function in individuals unable to exercise, preventing the metabolic decline that accelerates functional deterioration. Several next-generation AMPK activators are in development with improved tissue selectivity and potency compared to metformin. These include direct allosteric activators that bind specific AMPK subunits, as well as prodrugs optimized for oral bioavailability. Testing these compounds in preclinical models of age-related metabolic dysfunction and frailty should be a research priority.

### Therapeutic Implications and Translational Opportunities

Our findings suggest several immediately translatable interventions for sarcopenia prevention and treatment. First, exercise as molecular medicine, the complete reversal of MYORG and STRADB downregulation in active elderly individuals provides molecular validation for exercise as a primary prevention strategy. Guidelines already emphasize that even moderate regular activity (≥150 minutes/week) may prevent rather than merely slow sarcopenia development, with the mechanism running through preserved MYORG-mediated regenerative capacity and STRADB-mediated metabolic flexibility. Second, AMPK pharmacology through STRADB’s pivotal role in AMPK signaling identifies this pathway as a target beyond its existing diabetes indication. Although metformin’s effects on skeletal muscle exercise benefits are confounding, it has been suggested to have protective effects against sarcopenia and potentially improve sarcopenia related quality of life (Hu et al., 2025). Clinical trials testing newer selective AMPK activators specifically for sarcopenia prevention are warranted, particularly for individuals unable or unwilling to exercise. Third, preservation of regenerative capacity via MYORG’s function in myoblast differentiation suggests that interventions that support satellite cell activity, including growth factors, nutritional supplements, and pharmacological myogenic stimulators, may preserve youthful muscle turnover capacity. Monitoring MYORG expression could identify individuals at risk for regenerative failure before functional decline becomes clinically apparent. Fourth, precision medicine approaches: activity-dependent genes could stratify patients into those likely to respond to lifestyle interventions versus those requiring pharmacological support, while C4ORF54 and similar phenotype-mediated markers could track intervention efficacy in real time.

### Strengths and Limitations

This study’s major strength is the integrated discovery-validation design with systematic activity stratification. The discovery phase identified candidate genes in a well phenotyped cohort and grouped them by distinct functional categories. The validation phase tested these candidates across five independent studies spanning diverse activity contexts, enabling assessment of activity dependence. The consistency of findings across all five discovery genes, the negative correlation between activity controlled and sedentary datasets, and the poor validation of established atrophy markers all support the robustness of conclusions.

Several limitations warrant consideration. All datasets employed cross-sectional designs that compared young versus old adults, rather than following individuals longitudinally as activity changes. While the consistency across cohorts strengthens causal inference, longitudinal studies would provide definitive evidence. Such studies should track physical activity objectively (accelerometry), obtain serial muscle biopsies, and correlate gene expression changes with functional trajectories.

Physical activity assessment varied across datasets. GSE159217 used both questionnaires and accelerometry, but most datasets relied on protocols (e.g., 48-hour rest before biopsy) or sample characteristics (osteoarthritis patients with activity limitation) rather than direct activity measurement. More precise quantification of activity, including type, intensity, and duration, would enable dose-response analyses and the identification of optimal activity patterns for maintaining gene expression.

All datasets utilized the vastus lateralis muscle. While this consistency minimizes tissue-specific confounding, generalizability to other muscles requires verification. Different muscle groups exhibit distinct aging trajectories, proximal versus distal, antigravity versus non-antigravity, phasic versus tonic, potentially reflecting distinct activity patterns. Whether activity-protective effects are universal or muscle specific remains uncertain.

Validation focused on mRNA expression. While transcriptomics provides a comprehensive assessment and mRNA changes often correlate with protein changes, validating findings at the protein level would strengthen conclusions. Particularly for STRADB, demonstrating that mRNA changes translate to altered protein levels and AMPK activity would confirm functional relevance. Similarly, for MYORG, correlating expression with actual regenerative capacity (satellite cell function, repair after injury) would demonstrate phenotypic consequences.

Sample sizes in individual validation datasets were relatively modest (19-80 range in studies). While statistical power was adequate for detecting large activity effects, subtle effects may have been missed. Larger cohorts would enable more precise estimation of effect sizes and the identification of secondary patterns.

### Conclusions

This integrated discovery-validation study demonstrates that molecular signatures distinguishing fit from frail aging in skeletal muscle primarily reflect physical activity rather than inevitable biological aging. Discovery analysis revealed extensive bidirectional transcriptomic remodeling, including coordinated loss of muscle specific programs and active upregulation of RNA processing, chromatin remodeling, and stress response pathways. Systematic validation across activity-stratified cohorts showed that the top discovered genes, MYORG, STRADB, C4ORF54, LYPLA1, and OIP5-AS1, all exhibit activity-protective expression patterns maintained in active elderly but declining in sedentary aging.

The magnitude of activity effects (1.5-2.6-fold) matches or exceeds typical age effects, and the negative correlation between activity controlled and sedentary aging datasets (r = −0.55) demonstrates that genes respond in opposite directions depending on activity status. These findings fundamentally reframe skeletal muscle aging, revealing that what appears to be inevitable aging in conventional studies largely reflects preventable secondary aging driven by declining physical activity.

The activity protected pathways identified represent tractable therapeutic targets. MYORG-mediated regenerative capacity, STRADB-mediated metabolic flexibility, and C4ORF54-associated functional reserve all respond to mechanical and metabolic stimulation. Exercise remains the gold-standard intervention, but understanding activity responsive pathways enables the development of pharmacological approaches for individuals unable to exercise. AMPK activators, particularly metformin already FDA-approved for other indications, warrant immediate investigation for frailty prevention.

Longitudinal studies, exercise intervention trials with molecular endpoints, mechanistic investigations of activity responsive pathways, functional characterization of C4ORF54, and clinical trials testing AMPK activators will translate these molecular insights to tangible health benefits. The goal is not merely to extend lifespan but to extend healthspan, maintaining functional independence and quality of life into advanced age through science-based interventions targeting modifiable aging mechanisms.

## AUTHOR CONTRIBUTION

R.S. was responsible for the conceptualization, hypothesis generation, data analysis, compilation, review, editing, and figure generation. K.I. was responsible for conceptualization, review and editing.

## DECLARATION ON THE USE OF AI IN THE WRITING PROCESS

The authors declare that in the writing process of this work, no generative artificial intelligence (AI) or AI-assisted technologies were used to generate ideas or theories. AI technology, specifically Claude and Grammarly, was used solely to improve writing, enhance structure and readability, and refine language. This use was under strict oversight and control of and by the authors. The authors were responsible for the hypothesis, the interpretation of scientific findings, and the accuracy of the final content. All statements and conclusions reflect the author’s scientific judgment and are verified against primary literature. The authors fully comprehend that authorship comes with responsibilities and tasks that can only be attributed to and performed by humans, and the authors have adhered to these guidelines in the preparation of this manuscript.

## DATA AVAILABILITY STATEMENT

All datasets analyzed are publicly available through the Gene Expression Omnibus (GEO) database:

- Discovery: GSE144304
- Validation: GSE159217, GSE175495, GSE164471, GSE242202, GSE151066

Analysis code and processed data are available from the corresponding author upon reasonable request.

